# Phenotyping the virulence of SARS-CoV-2 variants in hamsters by digital pathology and machine learning

**DOI:** 10.1101/2023.08.01.551417

**Authors:** Gavin R Meehan, Vanessa Herder, Jay Allan, Xinyi Huang, Karen Kerr, Diogo Correa Mendonca, Georgios Ilia, Derek W Wright, Kyriaki Nomikou, Quan Gu, Sergi Molina Arias, Giuditta De Lorenzo, Vanessa Cowton, Nicole Upfold, Natasha Palmalux, Jonathan Brown, Wendy Barclay, Ana Da Silva Filipe, Wilhelm Furnon, Arvind H Patel, Massimo Palmarini

**Affiliations:** MRC-University of Glasgow Centre for Virus Research, United Kingdom; CVR-CRUSH, MRC-University of Glasgow Centre for Virus Research, United Kingdom; Department of Infectious Disease, Imperial College London, London, United Kingdom

**Author notes:** These authors contributed equally to this work. Deceased (Kyriaki Nomikou). Joint Senior Authors (Arvind H Patel, Massimo Palmarini).

## Abstract

SARS-CoV-2 has continued to evolve throughout the COVID-19 pandemic, giving rise to multiple variants of concern (VOCs) with different biological properties. As the pandemic progresses, it will be essential to test in near real time the potential of any new emerging variant to cause severe disease. BA.1 (Omicron) was shown to be attenuated compared to the previous VOCs like Delta, but it is possible that newly emerging variants may regain a virulent phenotype. Hamsters have been proven to be an exceedingly good model for SARS-CoV-2 pathogenesis. Here, we aimed to develop robust quantitative pipelines to assess the virulence of SARS-CoV-2 variants in hamsters. We used various approaches including RNAseq, RNA *in situ* hybridization, immunohistochemistry, and digital pathology, including software assisted whole section imaging and downstream automatic analyses enhanced by machine learning, to develop methods to assess and quantify virus-induced pulmonary lesions in an unbiased manner. Initially, we used Delta and Omicron to develop our experimental pipelines. We then assessed the virulence of recent Omicron sub-lineages including BA.5, XBB, BQ.1.18, BA.2 and BA.2.75. We show that in experimentally infected hamsters, accurate quantification of alveolar epithelial hyperplasia and macrophage infiltrates represent robust markers for assessing the extent of virus-induced pulmonary pathology, and hence virus virulence. In addition, using these pipelines, we could reveal how some Omicron sub-lineages (e.g., BA.2.75) have regained virulence compared to the original BA.1. Finally, to maximise the utility of the digital pathology pipelines reported in our study, we developed an online repository containing representative whole organ histopathology sections that can be visualised at variable magnifications (https://covid-atlas.cvr.gla.ac.uk). Overall, this pipeline can provide unbiased and invaluable data for rapidly assessing newly emerging variants and their potential to cause severe disease.

## INTRODUCTION

As the COVID-19 pandemic progressed over the past three years, the virus responsible for the disease, severe acute respiratory syndrome coronavirus 2 (SARS-CoV-2), has continued to evolve giving rise to a number of variants, some of which were defined as “variants of concern” (VOCs) by the WHO [1, 2]. These VOCs contain mutations, especially but not solely in the spike encoding S-gene, which may confer a selective advantage for example by increasing their transmissibility and/or immune evasion compared to the progenitor virus [1]. To date, the WHO has recognised five VOCs: B.1.1.7 (Alpha); B.1.351 (Beta); P.1 (Gamma); B.1.617.2 (Delta) and B.1.1.529 (Omicron; henceforth referred as BA.1 to differentiate it from other sub-lineages) [1, 3–5]. Since the emergence of the original BA.1 in November 2021 [5] different sub-lineages have emerged. Soon after the BA.1 emergence, BA.2 became the predominant variant followed by a variety of BA.2 descendants, including BA.5 and BA.2.75, which became predominant in some geographical regions [1, 6]. Although both BA.5 and BA.2.75 diversified from BA.2, these two Omicron sub-lineages are phylogenetically separated from each other, suggesting that BA.5 and BA.2.75 emerged independently. BQ.1.18 is a sub-lineage of BA.5 [7] while the XBB variant is a recombinant between BJ.1 (a BA.2.10 derivative) and BM.1.1.1 (a descendant of BA.2.75), which was first detected in September 2022 in India and spread significantly at the end of 2022 [8].

In general, each new variant spreading globally tends to be more transmissible than the previous dominant variant [1]. As population immunity increases, due to either vaccination or continuous virus exposure, it is likely that COVID-19 will adopt an endemic pattern, possibly with seasonal peaks, primed by variants evading pre-existing immunity in the population [9].

Understanding in “real-time” the degree of vaccine escape of any new variant is critical to determine vaccination policies. To this end, *in vitro* seroneutralisation assays have proven to be a useful surrogate to predict vaccine escape of SARS-CoV-2 variants. Virus virulence is another key phenotypic characteristic of any new variant requiring early assessment. The risk of severe disease and hospitalisation varies, with Alpha, Gamma and Delta VOCs carrying an increased risk of ICU admission compared to Beta and BA.1. Hence, the combination of increasing pre-existing immunity in the population together with the intrinsic attenuated characteristics of Omicron, has led to an overall decrease in the incidence of severe disease and mortality associated with COVID-19 [10–13].

It is however more difficult to predict the trajectory of new variants with respect to virus virulence. Although BA.1 has been shown to be attenuated, and assuming that its transmission potential is maintained, there are no universal evolutionary pressures that may keep this phenotypic trait in newly emerging sub-lineages or new VOCs.

We and others have shown that the observed reduction in virulence of the BA.1 variant correlates to a change in the virus entry pathways *in vitro*, and importantly in reduced virulence in experimentally infected hamsters [14–17]. Throughout the pandemic, small animal models have been used extensively to assess the virulence of wild type SARS-CoV-2 and emerging VOCs [8, 14, 16–26]. These studies have provided invaluable data on disease pathogenesis, virus transmission and the efficacy of different anti-viral compounds or vaccines [19, 27–32]. Importantly, hamsters have been shown to be naturally susceptible to SARS-CoV-2 infection and to be able to transmit the virus to humans [33]. In hamsters, BA.1 is unable to infect lung epithelial cells (unlike the original B.1 virus, Delta and other variants) [14, 16]. In addition, in experimentally infected hamsters it is also possible to recapitulate the increased virulence of the Delta variant compared to B.1 shown in the human population [23, 34].

Hence, although no animal model can fully recapitulate a human disease, hamsters represent an excellent model to dissect SARS-CoV-2 pathogenesis and determine the degree of virulence of newly emerging variants. To this end, many studies using experimentally infected hamsters, have focused on measuring *in vivo* viral replication, on identifying virus infected cells, and on examining pathogenic potential by measuring weight loss and assessing various histopathological criteria in general by qualitative scores [8, 17–20]. Here, we have developed unbiased and automated “digital pathology” methods to assess SARS-CoV-2 virulence. Digital pathology is a broad term that refers to a variety of systems to digitize pathology slides and associated meta-data, their storage, review, analysis, and enabling infrastructure [35]. Computational analysis of whole scanned tissue sections provides the opportunity to quantify cells or histological features in wide representative areas of infected organs. We applied these pipelines with recently evolved variants (BA.5, BQ.1.18, BXX, BA.2 and BA.2.75) [1, 6–8] and showed that some of them have gained a more virulent phenotype compared to the parent BA.1. This pipeline can contribute to the rapid assessment of newly emerging variants and should prove invaluable as the pandemic enters the next phase. Furthermore, we created an online repository to share with the scientific community high resolution digitized whole organ scanned slides from this study, providing a wider context to histopathology micrographs for experimental models of COVID-19.

## RESULTS

### Host transcriptional response to SARS-CoV-2 infection

To assess the complex host responses during SARS-CoV-2 infection, we first experimentally infected Golden Syrian hamsters with either the Delta (B.1.617.2) or the BA.1 variants. These variants are on the opposite spectrum of the phenotype associated with the clinical outcome of SARS-CoV-2 infection, both in humans and in experimental models. Hence, Delta and BA.1 can provide the baseline for the development of quantitative pipelines to assess virus virulence. As expected, between 2- and 6-days post-infection (dpi) the Delta-infected animals lost significantly more weight and had higher welfare scores than both the BA.1- and mock-infected hamsters (Fig. S1A), confirming the expected phenotype for both variants. We culled animals at either 2 or 6 dpi and collected tissues of both the upper and lower respiratory tract, in addition to peripheral blood, for downstream analyses.

To understand the overall host responses to SARS-CoV-2 infection and identify potential markers of virus virulence, we performed bulk RNAseq on lungs and blood of both infected and mock-infected hamsters. Principal component analysis indicated distinct clustering of both the Delta- and BA.1-infected groups at 2 dpi in lungs and blood but showed less distinctive separation of BA.1- and mock-infected animals at 6 dpi (Fig. S1B). Both in the lungs and in peripheral blood, Delta, and BA.1 induced significant differential gene expression compared to the mock-infected samples (Fig. 1A). As expected, both variants induced multiple cell responses pathways associated with antiviral mechanisms, immune system activation, cytokine and chemokine responses and interferon signalling as evident by gene ontology (GO) analysis (Fig. 1B, C). Direct GO analysis between the two variants found a limited number of pathways on day 6 upregulated only in delta-infected animals. These pathways include those associated with remodelling and lipoprotein regulation, suggesting that the differences in disease outcome caused by these two variants may be attributed to lesions and activation of tissue repair pathways in the lungs. Comparison of ∼200 interferon stimulated genes showed a general activation of the type-I IFN response in infected animals (as suggested by the GO analysis and previously published studies) both in the lungs and in the blood [36–38]. Delta-infected animals showed a more robust upregulation of ISGs than BA.1-infected hamsters in both lungs and blood (Fig. 1D, E). Overall, these analyses suggest that markers of type-I IFN response, immune system activation and tissue repair may be useful to characterise the extent of pathology induced by SARS-CoV-2 variants.

**Figure 1.**
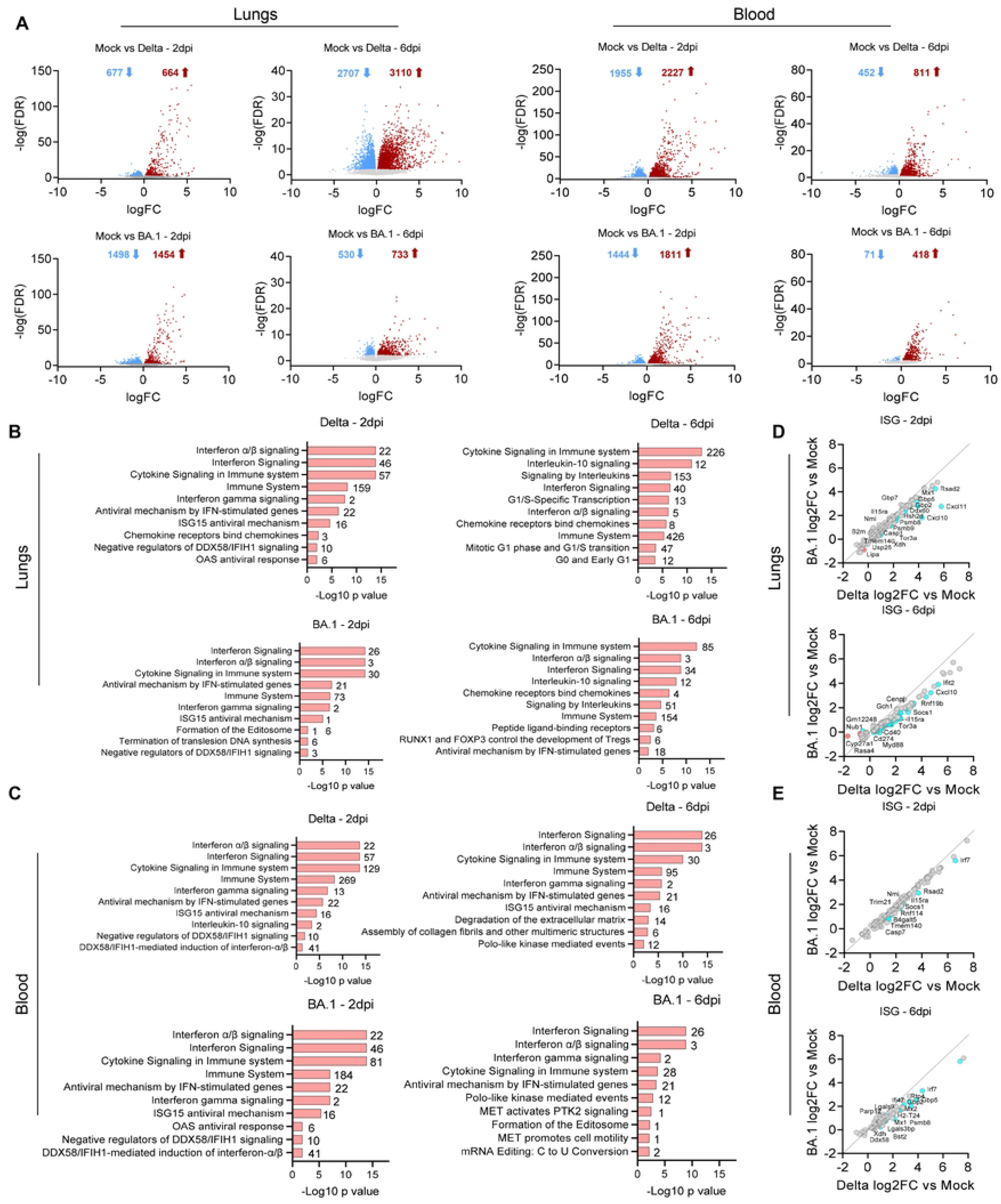
Transcriptomic response of lungs and blood of SARS-CoV-2 infected hamsters. (A) Volcano plots indicate the number of significantly upregulated (red arrows) or downregulated (blue arrows) genes in lungs or blood of hamsters infected with the indicated variant at each timepoint (compared to mock-infected controls). (B-C) Pathway analysis highlighting the enriched ontologies of differentially expressed genes in the lungs (B) and blood (C) of infected hamsters at 2 - and 6-days post-infection (dpi). The number of genes involved are indicated beside each pathway. (D-E) Scatterplots representing the relative expression of interferon stimulated genes (ISG) in lungs (D) and blood (E) of hamsters experimentally infected with either the Delta or BA.1 variant at the indicated timepoints. ISG with relative higher expression in BA.1 compared to Delta (FDR<0.05) are shown in red. ISG with relative higher expression in Delta compared to BA.1 (FDR<0.05) are shown in blue. Data derived from n=8 hamsters (4 females and 4 males per group), infected in two independent experiments.

### Imaging and quantification of SARS-CoV-2 replication in tissues

Next, we characterised the spread of SARS-CoV-2 infection in infected tissues. Throughout this study, for the detection of virus and cellular markers, we aimed to develop unbiased quantitative methods using software assisted whole section imaging and downstream automatic analyses including those enhanced by machine learning approaches (henceforth referred with the broad term of digital pathology) [39]. Also, in order to share the histopathology features shown in this study as comprehensively as possible, we have developed an online repository (“CVR Virtual Microscopy”; https://covid-atlas.cvr.gla.ac.uk) where whole organ scanned images can be accessed by users in their entirety and at variable magnification as if they were observing slides under a microscope.

First, we compared the detection levels of SARS-CoV-2 nucleocapsid protein with its spike RNA by immunohistochemistry (IHC) and *in situ* hybridisation, respectively. The two methods resulted in essentially identical staining patterns (Fig. S2). Background staining in mock-infected samples was higher using IHC and therefore we used RNA *in situ* hybridisation for the remaining part of the study. Next, we assessed virus replication for Delta and BA.1 along the entire respiratory tract. As expected, tissues collected from animals culled at 2 dpi showed both Delta- and BA.1-infected cells within the respiratory tract (Fig. 2A-B). We found instead little evidence of virus infected cells in animals culled at 6 dpi. At 2 dpi, we found no significant differences in the number of infected cells in the nose, larynx, and trachea of Delta- and BA.1-infected animals. In our samples, the nose represents the inner mucosa of the small cartilaginous tissue surrounding the nasal cavities of the hamster. However, both nasal turbinates of Delta-infected animals and the lungs showed a significantly higher number of infected cells than the same tissues in BA.1-infected hamsters. Importantly, significant spread of SARS-CoV-2 in the lung parenchyma was evident only in Delta-infected hamsters. Delta infected both epithelial cells in the bronchioles and alveoli forming large foci of infected tissues. Conversely, BA.1 infected only cells in the bronchioles and at most formed small foci of infection in the lung parenchyma in some animal (Fig. 2A-B). As expected, there was variability between animals within each group with respect to the number of infected cells. This variability was especially evident in the trachea, with 2 of 8 Delta-infected animals showing a number of infected cells more than 10-fold the number in the remaining animals of the group. Except for the two outliers indicated above, the tracheas of the remaining Delta- and BA.1-infected hamsters showed a similar number of infected cells.

**Figure 2.**
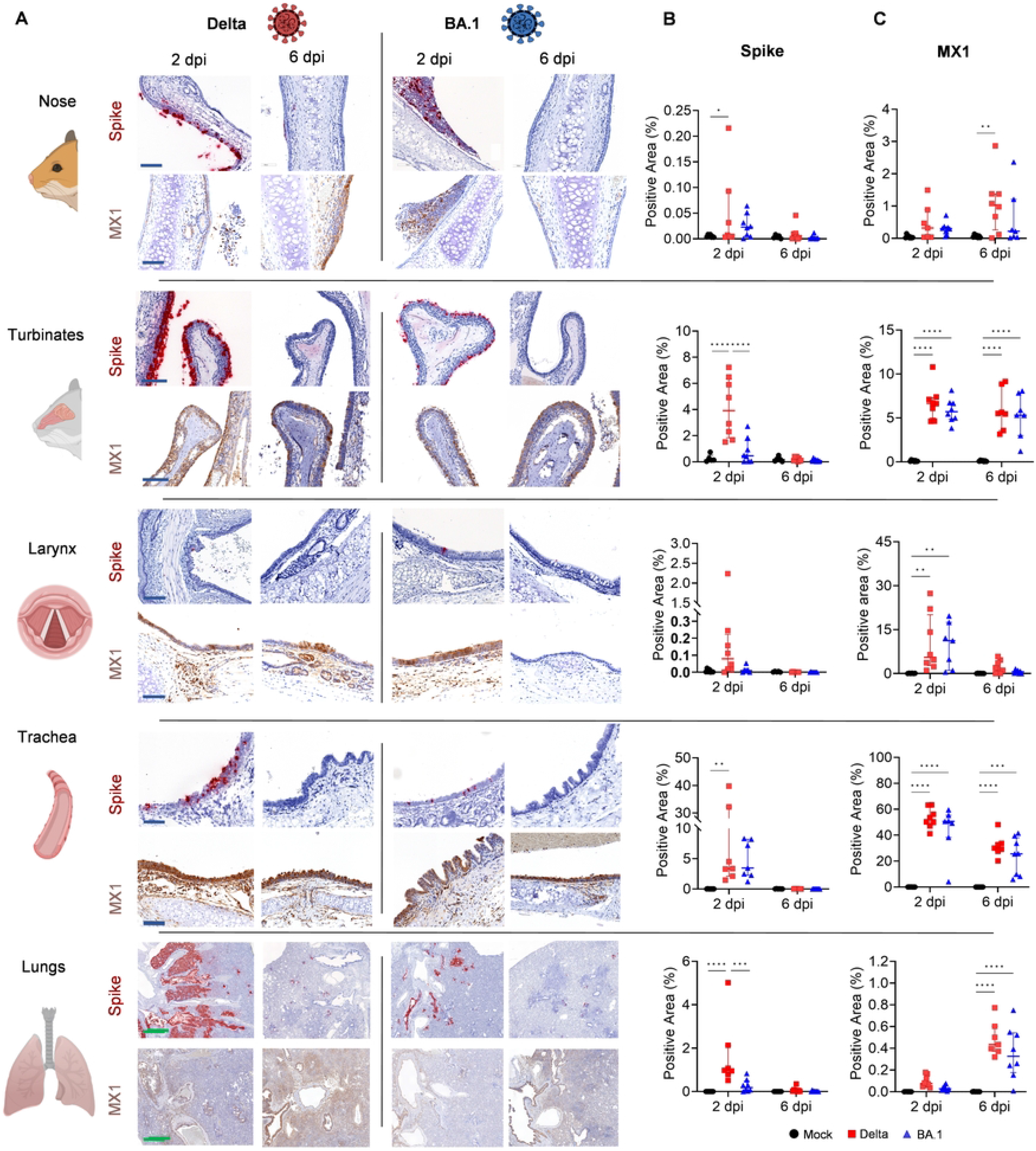
Distribution of Delta and BA.1 in organs of the respiratory tract of experimentally infected hamsters. Golden Syrian hamsters were infected intranasally with either Delta or BA.1 (or mock-infected). Control animals received media alone. Animals were culled 2- or 6-days post infection (dpi) and the nose, turbinates, larynx, trachea, and lungs were collected for digital pathology analyses (A). Tissues were assessed for the presence of spike RNA by *in situ* hybridisation (B) or for the expression of MX1 by immunohistochemistry (C). For signal quantification, slides were scanned with an Aperio VERSA 8 Brightfield, Fluorescence & FISH Digital Pathology Scanner (Leica Biosystems) at 200 x brightfield magnification. Areas to be analysed were outlined using QuPath (Version 0.3.2), HALO® (Indica Labs) or Aperio ImageScope (Leica Biosystems). The algorithm to detect the percentage of positively stained area in pixel was tuned individually before analysis. Statistical analysis was performed using a Two-Way ANOVA, *<0.05, **<0.01, ***<0.001, ****<0.0001. Data were derived from two independent experiments (n=8 in total; n=4 per experiment). Black circles: uninfected; red squares: Delta-infected; blue triangles: BA.1-infected. Blue scale bar: 100 μm; green scale bar: 1 mm. Graphics made using biorender.com.

Given the data obtained by RNAseq, where the interferon response is a key differentially activated pathway in infected hamsters, we also assessed expression of MX1, as a representative core interferon stimulated gene [40]. MX1 expression in the nose and lungs was in general lower at 2 dpi compared to the levels observed at 6 dpi. The larynx showed high MX1 expression on 2 dpi but little on 6 dpi, while the turbinates and trachea showed similar levels at both timepoints (Fig. 2C). Bulk RNAseq data suggested a more robust type-I IFN response elicited by the Delta variant compared to BA.1 in lungs from infected hamsters (Fig. 1B-E). We aimed therefore to spatially resolve ISG expression in infected hamsters using *in situ* RNA hybridisation of serial sections of tissue lungs. We used five sections with the middle section probed for spike, while in the other sections we used RNA probes for the following ISGs: RSAD2, IFIT1, MX1 and OAS1 (Fig. 3). The number of ISG-positive cells was directly related to the number of spike-positive cells in the lungs of infected hamsters. There were clear overlapping virus- and ISG-positive areas in both Delta- and BA.1-infected lungs. Hence, the more robust type-I IFN response observed by RNAseq in the lungs and blood of infected animals correlates with higher replication levels of Delta in the lungs.

**Figure 3.**
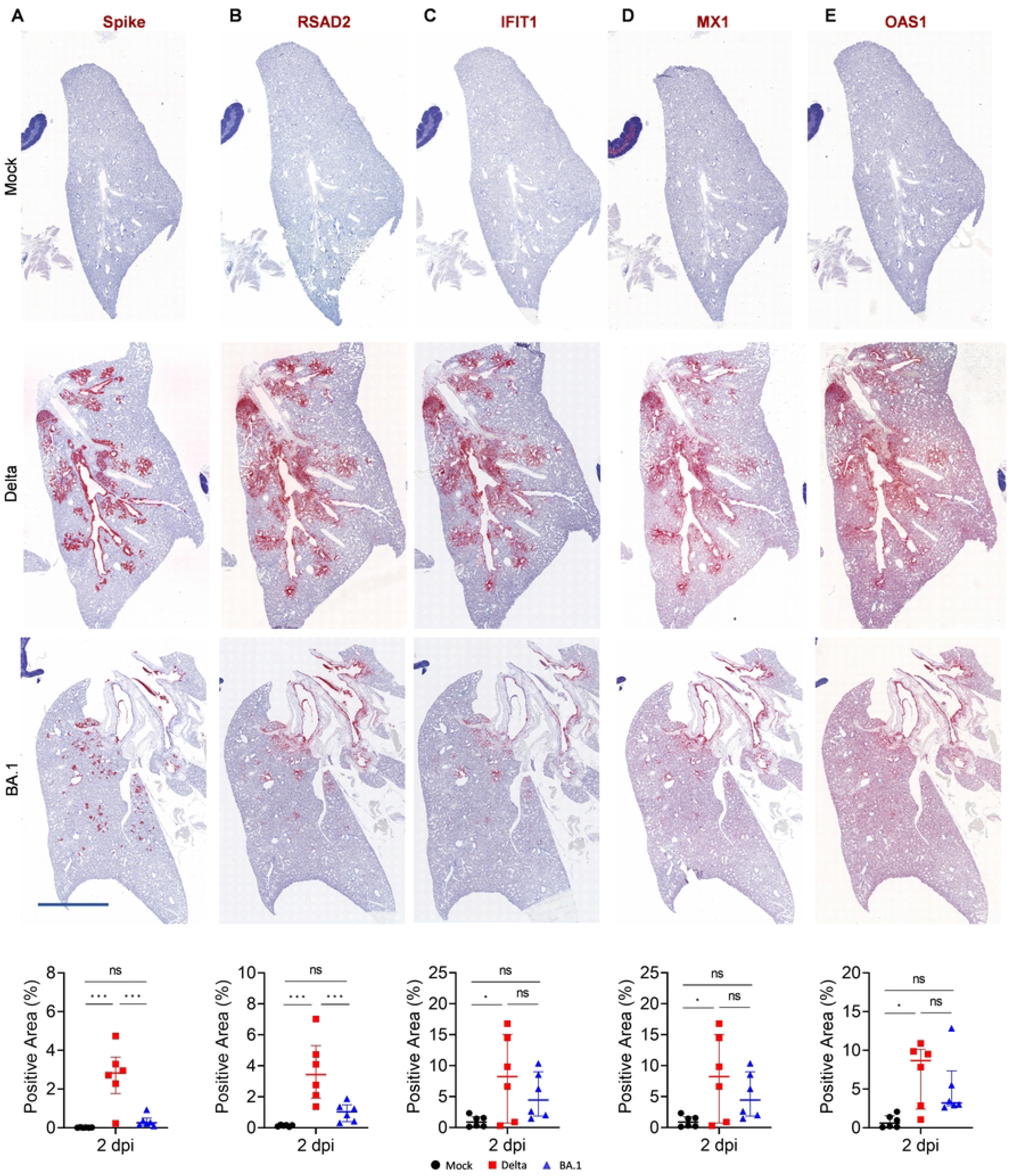
Expression of interferon stimulated genes in lungs of experimentally infected hamsters. (A) *In situ h*ybridisation of serial lung sections obtained from hamsters experimentally infected with either Delta or BA.1 and killed at 2 days post-infection (2 dpi). Probes used included those for SARS-CoV-2 spike, RSAD2, IFIT1, MX1 and OAS1. (B) Signal was quantified as in Fig. 2 and statistical analysis was performed using a One-Way ANOVA, *<0.05, **<0.01, ***<0.001, ****<0.0001. Data represents two independent experiments using a total of n=6 hamsters (3 females and 3 males per group). Black circles: uninfected; red squares: SARS-CoV-2 (Delta) infected; blue triangles: SARS-CoV-2 (BA.1) infected. Scale bar: 3 mm.

The differences between Delta- and BA.1-infected hamsters were also present at earlier timepoints. Analysis of tissues collected at 1 dpi of the turbinates, trachea, and lungs (Fig. S3A) showed increased levels of spike RNA (Fig. S3B) and MX1 expression (Fig. S3C) in Delta-infected animals. Similarly, nasal washes, throat swabs and whole lung RT-qPCR indicated higher levels of genomic RNA in Delta-infected hamsters (Fig. S3D).

### Quantifying the extent of pulmonary lesions by digital pathology

We showed above that at 6 dpi, at the peak of clinical signs in experimentally infected hamsters, most of SARS-CoV-2 has been cleared by the host (Fig. 2A). Our RNAseq analysis suggested that in addition to markers of the type-I IFN response, pathways leading to immune cell activation and tissue repair are also differentially upregulated between Delta- and BA.1-infected hamsters (Fig. 1B-C). Histopathology lesions in the lungs of infected hamsters (especially those infected with Delta) were characterised by infiltration of macrophages in the alveoli and in the interstitium with a multifocal to coalescent distribution especially at 6 dpi (Fig. 4). The immune cell infiltration also contained neutrophils/heterophils as well as lymphocytes and plasma cells. At 2 dpi, Delta-infected hamsters showed vasculitis and a sloughing of bronchial epithelial cells. Vascular and bronchial lesions was instead minimal in BA.1-infected animals at 2 dpi. As shown in other studies [29, 41, 42], we found marked alveolar epithelial hyperplasia forming dense meander- and rosette-like structures replacing the alveolar spaces especially in the Delta-infected hamsters (Fig. 4). We termed these lesions “medusa,” as they resemble the shapes of the venomous snakes of the mythological Greek Medusa. The cellular infiltrates and the proliferation of the type 2 pneumocytes (or other lung progenitor cells), which starts around the bronchi involved a large area of the lung (around 50%), especially in Delta-infected hamsters. Occasionally, multinucleated cells (interpreted as syncytia) were associated with the often severely hyperplastic bronchial epithelium or in the lung parenchyma (Fig. 4).

**Figure 4.**
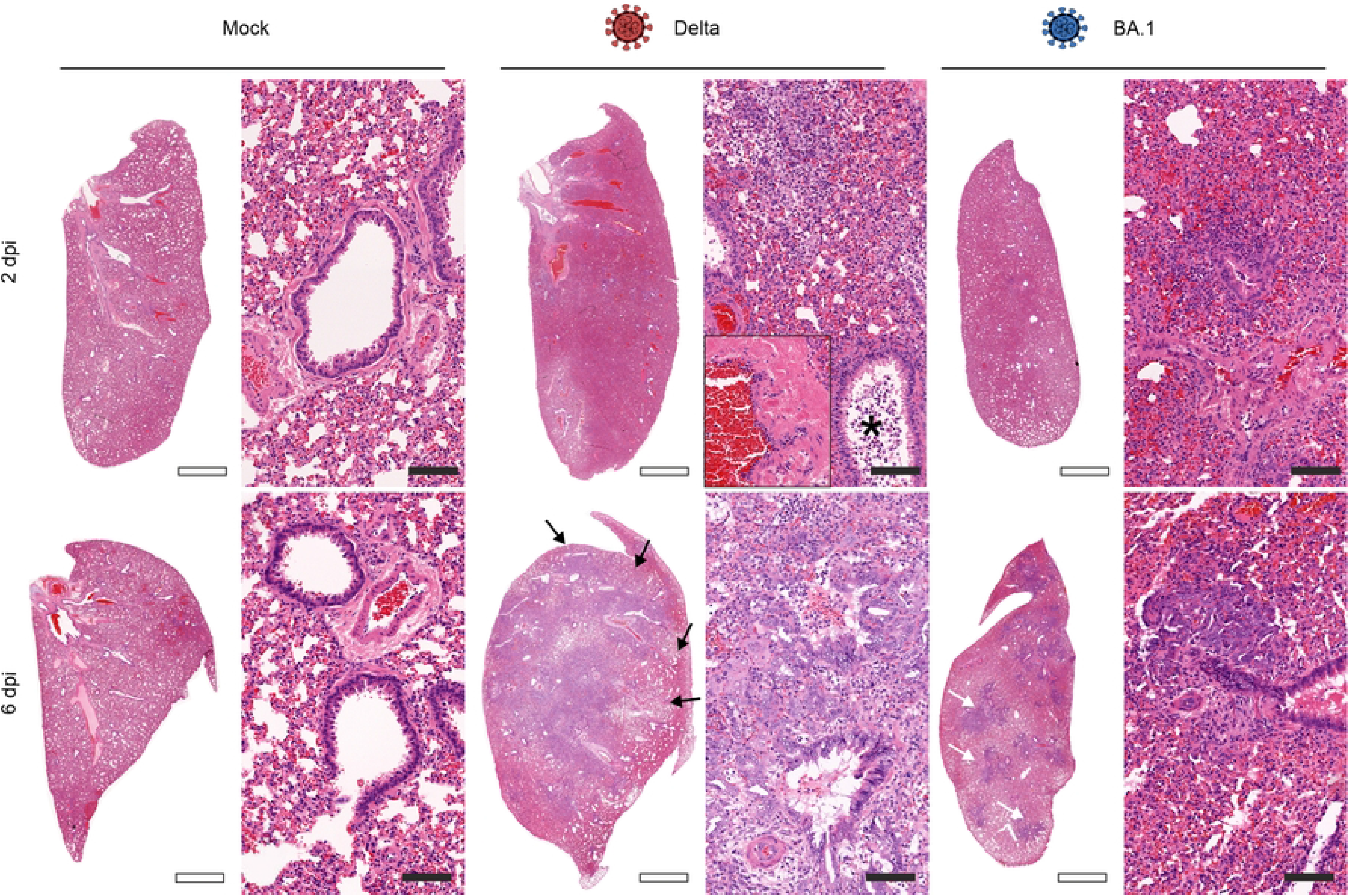
Histopathology of lungs of experimentally infected hamsters. Low (left) and high (right) magnification of lung sections stained with haematoxylin and eosin of hamsters experimentally infected with either the Delta or BA.1 variant (or mock-infected controls). At 2 days post-infection (dpi), Delta causes a higher level of infiltration of macrophages in and around bronchi, while the BA.1 variant causes only mild infiltrations at 2 dpi compared to mock-treated hamsters. On the same day, Delta-infected animals showed a moderate vasculitis (inset, right panel) and a moderate sloughing of bronchial epithelial cells (asterisk). Vascular and bronchi pathology was minimal in BA.1-infected animals at 2 dpi. At 6 dpi the lungs of Delta-infected animals show a severe infiltration of macrophages, lymphocytes, plasma cells and neutrophils/heterophils as well as a severe alveolar epithelial hyperplasia covering large areas of the lung lobe (black arrows). BA.1-infected animals show in some cases the same lesions but covering only limited amounts of the lung lobes (white arrows). Empty scale bars: 2 mm; filled scaled bars: 100 μm.

Hence, we next aimed to image and quantify the extent of virus-induced pulmonary lesions by first evaluating and comparing the immune cells infiltrate in the lungs of Delta- and BA.1-infected hamsters. We specifically assessed T cells (CD3^+^) and macrophages (IBA1^+^) and found a significant increase in the number of these cells in Delta-infected hamsters compared to those infected with BA.1 (Fig. 5A-B. By histopathology, both cell types represent most immune cell infiltrates in the lungs of Delta-infected hamsters (Fig. 5A, B).

**Figure 5.**
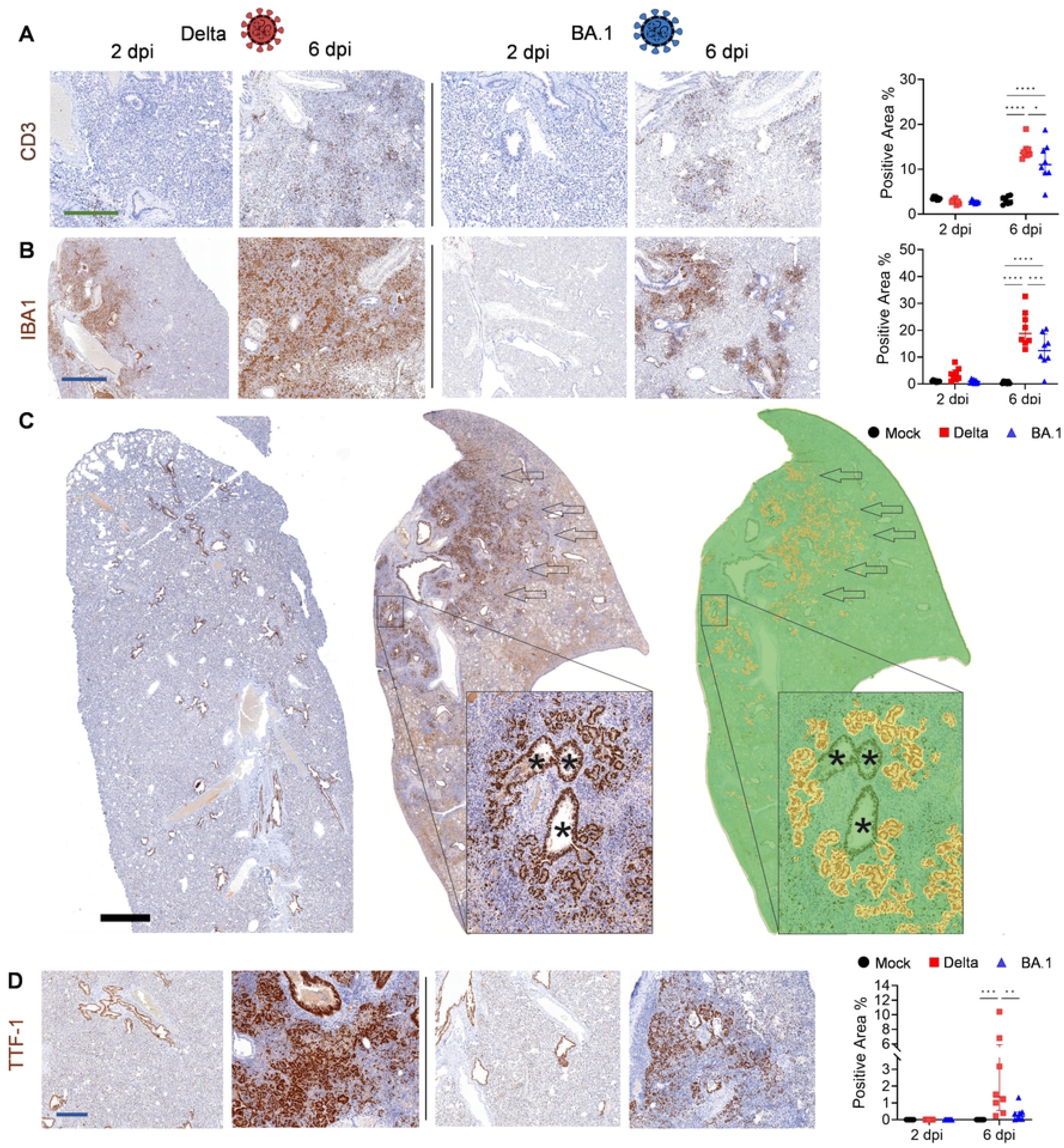
Quantification of tissue histopathology in Delta or BA.1-experimentally infected hamsters. Images of whole lung sections of hamsters experimentally infected with either Delta or BA.1 (or mock-infected controls) and culled at either 2- or 6-days post-infection (dpi). Expression of CD3 (A), IBA1 (B) and TTF1 (C-D) was assessed by *in situ* hybridisation. Animals were culled 2- or 6-days post infection (dpi) and the lungs were collected for histological analysis. Note that TTF1 is expressed by hyperproliferating epithelial cells (arrows), isolated type-2 pneumocytes and by epithelial cells lining the bronchi/bronchioles (indicated with an asterisk). The software HALO was therefore trained to capture only proliferating alveolar epithelial cells (“medusa”) and exclude isolated TTF1^+^ type-2 pneumocytes and bronchial cells. CD3- and IBA1-positive cells in experimentally infected animals were quantified using QuPath (Version 0.3.2), while HALO was used to quantify medusa. Statistical analysis was performed using a Two-Way ANOVA, *<0.05, **<0.01, ***<0.001, ****<0.0001. Data were obtained from two independent experiments using 8 hamsters (4 females and 4 males infected in two independent experiments). Black circles: uninfected; red squares: SARS-CoV-2 (Delta)-infected; blue triangles: SARS-CoV-2 (BA.1)-infected. Blue scale bar: 1 mm; green scale bar: 400 μm. Graphics made using biorender.com.

We next developed a method to quantify alveolar epithelial hyperplasia. We used the thyroid transcription factor (TTF1) [43, 44], a critical factor required for the expression of the surfactant protein in the respiratory epithelia. As expected, we found that TTF1-positive cells included the hyperplastic alveolar epithelial cells but also normal/isolated type 2 pneumocytes in the lungs and epithelial cells in the terminal bronchioles. To quantify only the hyperplastic areas, we used software assisted imaging detection. Using supervised machine learning approaches, we “trained” the HALO software (Indica Labs) to detect clusters of TTF1-positive nuclear areas representing proliferating type-2-pneumocytes while ignoring isolated type 2 pneumocytes or TTF1^+^ cells in the terminal bronchioli. We found no hyperplastic lesions at 2 dpi in any of the hamster groups while at day 6 we found significantly more medusas in Delta-infected hamsters compared to those infected with BA.1 (the latter had only values just above background in most animals) (Fig. 5C, D).

### Assessing the virulence of Omicron sub-lineages

The pipelines developed so far allow us to provide an automatic, unbiased, and semi-quantitative method to assess the degree of virulence of SARS-CoV-2 in hamsters. Hence, we next used this method to assess virulence of recently emerged omicron sub-lineages.

We first investigated BA.5, as other studies, although carried out with chimeric BA.2/BA.5 viruses, suggested that this variant had an increased virulence compared to BA.1 [22] (Fig. 6A, B). Lungs collected from hamsters infected with BA.5 showed, in comparison to those infected with BA.1, an increase in (i) macrophage infiltrate (IBA-1^+^ cells), (ii) cells expressing MX-1 and (iii) alveolar epithelial hyperplasia, medusas (Fig. 6A). These changes were however not as pronounced as those in the lungs of Delta-infected animals and did not reach statistical significance compared to values in BA.1-infected hamsters. BA.5-infected animals also showed a small decrease in body weight at 6 dpi, while animals infected with Delta or BA.1 showed the expected phenotype (slight increase in weight for BA.1-infected animals and weight loss for Delta from 2 dpi) (Fig. 6B).

**Figure 6.**
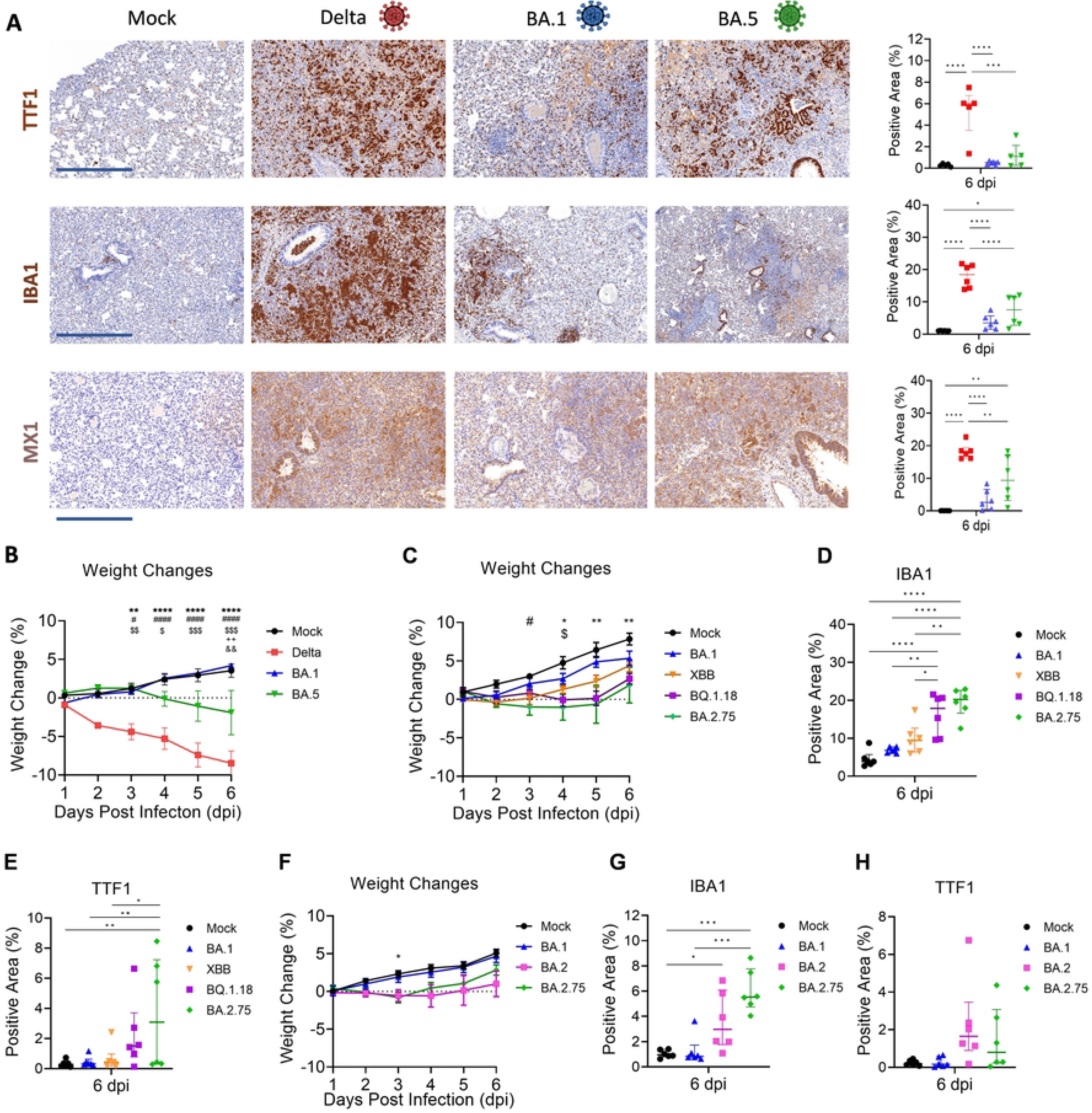
Quantification of Omicron sub-lineages virulence in experimentally infected hamsters. (A) Micrographs of lung sections collected at 6 days post-infection (dpi) from hamsters experimentally infected with either Delta, BA.1, BA.5 or mock-infected. Expression of TTF1^+^ hyperplastic epithelial cells and infiltrating macrophages (IBA1^+^) was assessed by *in* situ hybridisation. Expression of MX1 was instead assessed by immunohistochemistry. Data were quantified as in Fig. 5. (B) Differences in the weights between these animals were recorded daily. (C) Daily recorded weight changes in animals mock-infected or infected with either BA.1, XBB, BQ.1.18 or BA.2.75. Lung whole scanned sections were analysed for the presence of IBA1^+^ cells (D) or hyperplastic alveolar epithelial cells (E) as in A. (F) Daily weight changes of mock-infected hamsters or animals infected with either BA.1, BA.2 or BA.2.75. Quantification of cells expressing IBA1 (G) or hyperplastic TTF-1^+^ cells was conducted as in A. Statistical analysis was performed using a One-Way ANOVA with Tukey’s multiple comparisons test. Weight comparisons were performed using a Two-Way ANOVA. Significance is indicated with *<0.05, **<0.01, ***<0.001, ****<0.0001. * Denote comparisons between Delta and mock or BQ.1.18 and mock or BA.2.75 and mock; # denote comparisons between BA.1 and delta or mock and BA.2.75; $ denotes comparisons between Delta and BA.5 or mock and XBB; + denotes comparisons between uninfected and BA.5; & denotes comparisons between BA.5 and BA.1. n=6 (3 females and 3 males per group). Black circles: uninfected; red squares: SARS-CoV-2 (Delta) infected; blue triangles: SARS-CoV-2 (BA.1) infected; green inverted triangles: SARS-CoV-2 (BA.5) infected; orange inverted triangles: SARS-CoV-2 (XBB) infected; purple squares: SARS-CoV-2 (BQ.1.18) infected; pink squares: SARS-CoV-2 (BA.2) infected; green diamonds: SARS-CoV-2 (BA.2.75) infected. Scale bar: 500 μm. Graphics made using biorender.com.

Given that the digital pathology pipeline was able to show a possible intermediate virulence phenotype for BA.5, we proceeded to assess the virulence of other Omicron sub-lineages such as BQ.1.18, XBB and BA.2.75 (Fig. 6C-E). None of the variants induced major weight loss in infected hamsters (Fig. 6C). Hamsters infected with BA.2.75 showed a modest weight loss between dpi 2 and 5 but neither hamsters infected with BA.2.75 nor BQ.1.18 show, unlike animals infected with BA.1, weight increase until 5 dpi (Fig. 6C).

We found an increase in macrophages infiltrating the lungs in hamsters infected with the other variants compared to those infected with the original BA.1 (Fig. 6D). BA.2.75 showed the highest levels of IBA^+^ cells in the lungs, although differences were statistically significant only with BA.1 and XBB, but not with BQ.1.18. Lungs of animals infected with BA.2.75 also displayed a significant increase of alveolar epithelial hyperplasia compared to lungs of BA.1-infected animals (Fig. 6E). Overall, the data suggest that virulence of Omicron sub-lineages, and especially BA.5 and BA.2.75 increased compared to BA.1. We also compared BA.2.75 to its predecessor BA.2 to determine whether there had been further evolutionary adaptations in BA.2.75 that favoured a more virulent phenotype. No significant differences were observed for weight losses and alveolar epithelial proliferation between BA.2.75 and BA.2 (Fig. 6F-G). However, animals infected with BA.2.75 or those infected with BA.2 did not gain weight as steadily as mock- or BA.1-infected animals. We saw however a trend for BA.2.75-infected hamsters to show an increase in the levels of infiltrating macrophages in the lungs compared to BA.2, although differences were not statistically significant (Fig. 6H). Finally, to visualise and compare all the different variants used in this study, we normalised the data using values obtained in BA.1-infected hamsters as unit of reference between different experiments (Fig. S4). Graphs displayed in Fig. S4 show, as expected, a gradient of virulence between variants. XBB induced little or no lung pathology similarly to BA.1. BA.5, BA.2 and BQ.1.18 were more virulent than BA.1, while BA.2.75 was clearly more virulent than BA.1 but not as virulent as Delta. The use of a standard virus of reference could therefore enable comparisons between variants used in different experiments (and different laboratories).

## Discussion

The COVID-19 pandemic has entered a phase characterised by the periodic emergence of immune escape variants, which may differ in their virulence from their predecessors. Assessing the virulence of any newly emerged variant will be therefore one of the key features requiring near real-time monitoring. Animal models have been used throughout the pandemic to unveil many aspects of SARS-CoV-2 pathogenesis [42]. So far, there has been an excellent correlation between the virulence of SARS-CoV-2 variants such as Delta and BA.1 in humans and in experimentally infected hamsters [7, 17–20, 25, 26, 41, 45, 46].

In many studies, virulence of SARS-CoV-2 variants in experimentally infected hamsters has been determined by assessing body weight loss, and various features of lung function and histopathological lesions [8, 14, 16, 17, 22–26, 47]. In general, these parameters have proven to be good proxies of the virulence of variants. This is particularly so for variants such as Delta and Omicron, which exhibit clearly distinct phenotypes with respect to virulence. Some relative discrepancies between studies can, however, arise when the intrinsic differences in virulence between variants are modest, as for example with BA.2 and BA.5 compared to BA.1 [22, 25].

Body weight and pulmonary function could be theoretically affected by a variety of factors related to animal and welfare management that may differ in different experimental settings. However, histopathological features in infected animals such as alveolar damage and inflammatory lesions in the respiratory tract are a consequence of virus replication (and host immune responses) and therefore directly reflect virus virulence. In various studies, lung pathology is often characterised by qualitative scores determined by trained pathologists on lesions such as bronchiolitis, haemorrhages, alveolar damage and others. Qualitative histopathology scores can vary between individuals, and therefore these types of data are difficult to compare unequivocally between different laboratories. In this study, we aimed to develop a framework for quantitative unbiased methods to phenotype the relative virulence of SARS-CoV-2 variants by quantifying the extent of pulmonary pathology in experimentally infected hamsters. In line with previous studies [38, 48–50], our transcriptomic analysis showed that SARS-CoV-2 infection induces an infiltrate of immune cells in the lungs and immune activation in general. Indeed, here we found that expression of ISGs, and infiltrate of T cells and macrophages are common in lesions induced by the hypervirulent variant Delta but are barely above background in BA.1-infected hamsters.

In addition, our RNAseq data suggest that tissue remodelling was another key parameter distinguishing the transcriptional profile of Delta- and BA.1-infected hamsters. Virulent SARS-CoV-2 variants induce alveolar damage, with necrosis of type 1 and type 2 pneumocytes. Lung damage is subsequently repaired by alveolar epithelial hyperplasia, which is a key feature of lesions induced by SARS-CoV-2 in experimentally infected hamsters [29, 41, 42], and it is also found in some cases in post-mortem samples of human patients dying as result of COVID-19 [51–53]. Lung respiratory epithelium repair is due to proliferation of either type 2 pneumocytes following mild injuries [54], or other lung progenitor cells in response to severe injury with abundant loss of type 1 pneumocytes [55–57].

Overall, we found that infiltrating macrophages and proliferating epithelial cells constitute most of the cells in the affected areas of the lung parenchyma of SARS-CoV-2-infected hamsters. These two features represent therefore exceedingly good markers that by themselves provide an unbiased quantification of the virus-induced pulmonary lesions, and by extension virus virulence. In addition, the use of whole lung sections can also reduce bias by providing a spatial overview of the inflammatory response. For example, our investigation of the spatial distribution of the ISG response and how it directly correlated to the presence of virus in an inflammatory lesion proved to be particularly insightful (Fig. 3). Whole scanned imaging of tissue sections and downstream analyses including those based on artificial intelligence approaches (“digital pathology”) have been used extensively in diagnostic pathology and other fields including cancer research and infectious diseases [58–60].

In our study, by performing immunohistochemistry of whole scanned lungs sections to identify IBA1^+^ cell (as convenient marker for macrophages), we were able to quantify infiltrating macrophages in the lung parenchyma of experimentally infected hamsters. In this manner, the relative number of infiltrating macrophages in the lung parenchyma can be acquired in an unbiased fashion. The same approach to measure alveolar epithelial hyperplasia by simply quantifying TTF1^+^ cells did not initially provide satisfactory results, due to the expression of this marker in both proliferating and non-proliferating type 2 pneumocytes, as well as bronchial epithelial cells. However, supervised machine learning approaches allowed us to train the software used to detect clusters of TTF1^+^ cells (histological structures that we termed “medusa”) and exclude isolated type 2 pneumocytes (representing the normal type 2 cells in the lungs) or TTF1^+^ cells in the bronchiolar epithelium. Our approach allowed us not only to clearly distinguish the pulmonary lesions between those induced by the virulent Delta and the attenuated BA.1, but also to show an increased virulence of other Omicron sub-lineages.

The pipeline used in this study, enabled us to show that BA.5 and BA.2.75 have acquired an increased virulence phenotype compared to BA.1, as also suggested in other recent studies [6, 22, 26]. We saw no differences instead in virulence between XBB and BA.1, also in keeping with another recent study [8]. Interestingly we found a tendency for BA.2 to induce a higher number of macrophage infiltrates and medusas compared to BA.1, although differences did not reach statistical significance. Other published studies found no major differences in virulence between BA.1 and BA.2 [8, 47], in contrast to a previous study which used recombinant viruses with either BA.1 or BA.2 spike [61].

We propose that the approach described in this study can be used to quantify moderate differences in virulence between variants, although animal group sizes may need to be adjusted to address specific experimental questions. Indeed, most studies focusing on SARS-CoV-2 virulence in hamsters use experimental groups between 4 and 6 animals (and often males only), but these may not be sufficient to reveal modest differences in virulence between variants. It may be argued that modest differences in variant virulence observed in hamsters could have limited biological significance in human patients. Indeed, a limitation of these type of studies, is that the intrinsic virulence of variants in naïve hamsters is difficult to compare with their virulence in the “real world” in the human population, with pre-existing immunity derived from vaccination or previous infections. However, the intrinsic virulence of newly emerging variants remains a key phenotype to monitor to determine whether adjustments to public health measures are needed. For example, the emergence of a variant with increased virulence may require different vaccination policies from the existing ones.

We use equal number of both male and female hamsters in all our experiments. As expected, we observed some gender-independent variability in the extent of the lesions caused by the same variant between individual hamsters both within and between different experiments. A standard reference virus could be used in multiple experiments assessing different variants to normalise the data between experiments (Fig. S4). This approach would also enable meaningful comparison of data between different laboratories.

In conclusion, the digital pathology pipelines developed in this study provide a framework for the quantitative assessment of virulence of SARS-CoV-2 variants in experimentally infected hamsters. The identification of virus standards with defined pathogenicity and sharing of protocols for the quantification of pulmonary lesions, can allow comprehensive comparison of *in vivo* data between different laboratories. We also developed an online repository to share more effectively with the research community histopathological images derived from scanned whole organ tissue sections. We argue that the reader will be able to appreciate better histopathological micrographs contained in the main body of manuscripts if these were supplemented by images of whole scanned slides as those contained in our virtual microscope. Due to space constraints, micrographs contained in figures of standard manuscripts show only individual areas of interest at single or two magnifications. Virtual microscopes allow the reader to view all areas of a given section and at multiple magnifications, providing therefore a comprehensive spatial context of the data.

## MATERIALS AND METHODS

### Cells and Viruses

Calu-3 cells (ATCC HTB-55) are human lung adenocarcinoma epithelial cells. African green monkey kidney cells (Vero E6) expressing the human Ace2 receptor [62] were used to propagate the viruses. All cell lines were maintained at 37°C and 5% CO_2_ in DMEM (ThermoFisher) supplemented with 10% foetal bovine serum (FBS) (ThermoFisher), except for Calu-3 cells where RPMI-1640 medium (ThermoFisher) was supplemented with 20% FBS. BA.5 virus isolate was obtained as passage P2 from Greg Towers (University College London, UK) after initial access to passage P1 obtained from Alex Sigal (AHRI, South Africa). Other variants used in this study included the Delta variant (B.1.617.2, GISAID accession number EPI_ISL_1731019), BA.1, BA.2, BA.2.75, BQ.1.18, and XBB all isolated from clinical samples obtained either at the CVR or at the Imperial College London. Viruses were isolated from a clinical nasopharyngeal swab sample collected in virus transport medium. Samples were then resuspended in serum-free DMEM supplemented with 100 units mL-1 penicillin-streptomycin, 10 ug mL-1 gentamicin and 2.5 ug mL-1 amphotericin B to a final volume of 1.5 mL. Calu-3 cells were then inoculated and incubated overnight at 37C, 5% CO2. Next day, cell culture medium was replaced with fresh complete medium before further incubation. Infected cells were monitored for signs of CPE and the presence of viral progeny in supernatant by RT-qPCR. All working stocks were propagated in VeroE6 cells.

### Animals

Golden Syrian hamsters (HsdHan®:AURA were bred and maintained by Envigo; Wyton, United Kingdom). Animals were shipped as required and acclimatised at the Veterinary Research Facility (VRF) at the University of Glasgow prior to transfer to Containment Level 3 (CL3) suite. Animals were maintained in individually ventilated cages (IVCs) on a 12-hour light/dark cycle and provided with food and water *ad libitum*. Both male and female hamsters between 8-12 weeks old were used for experiments. All procedures were performed under a UK Home Office licence in accordance with the Animals (Scientific Procedures) Act 1986. All animal research adhered to ARRIVE guidelines.

### Experimental infections

Animals were randomised to treatment groups, however, due to the nature of this work blinding was not possible. Animals were handled within a Class I microbiological safety cabinet (MSC) in a CL3 suite throughout the experiment. Hamsters were anaesthetised with oxygen (1.5 L/minute) containing 5% isoflurane and intranasally dosed with 50 μl of Dulbecco’s Modified Eagle Medium (DMEM) containing 3.75×10^6^ genome copies (the equivalent of approximately 1×10^4^ PFU) of SARS-CoV-2. We chose to use genome copies for our infection studies to control for the differences in infectivity and plaque formation that have been observed with the BA.1 variant [15]. Control animals received DMEM only. Animal weights and temperatures were recorded daily, and animals were monitored twice daily for signs of clinical disease including piloerection, hunching, abnormal breathing and reduced peer interactions. Animals received a disease score based upon weight loss and the presence and severity of these signs. Animals were culled at the end of the experiment via a rising concentration of CO_2_.

### Throat Swabs

Animals were restrained and swabs (MWE) were inserted into the mouth and rotated five times on each tonsil. The swabs were then placed in 2 ml DMEM (ThermoFisher) containing 2% FBS (ThermoFisher) for 1 minute. For RNA RT-qPCR, 250 μl of media was added to 750μl TRIzol LS (ThermoFisher). For live virus assays, the media was frozen at - 80°C.

### RNA extractions

Viral RNA was extracted from culture supernatants using the RNAdvance blood kit (Beckman Coulter Life Sciences) following the manufacturer’s instructions. Tissue samples, approximately 20 mg in size, were collected in homogenisation tubes containing 2.8 mm metal beads (Stretton Scientific) and 1 ml TRIzol reagent (ThermoFisher Scientific). Blood was collected in TRIzol LS reagent (ThermoFisher Scientific) at a 1:4 ratio. Tissue samples were homogenised (6500 rpm, 4x 30s cycle, 30s break, room temperature) using a Precellys Evolution Homogeniser (Bertin Instruments) and the homogenate was mixed with 200 μl chloroform (Merck - 1L). Each sample was mixed thoroughly and centrifuged at 12,000 x g for 15 minutes at 4°C. The aqueous phase was removed and mixed with 230 μl 100% ethanol (Merck). The samples were then added to columns from RNeasy Mini Kits (Qiagen) and processed as per manufacturer’s instructions. On column DNase digestion (Qiagen) was performed on all samples for 15 minutes as per manufacturer’s instructions.

### RT-qPCR

RNA was used as template to detect and quantify viral genomes by duplex reverse transcriptase (RT) quantitative polymerase chain reaction (RT-qPCR) using a Luna Universal Probe one-step RT-qPCR kit (New England Biolabs, E3006E). SARS-CoV-2-specific RNAs were detected by targeting the ORF1ab gene using the following set of primers and probes: SARS-CoV-2_Orf1ab_Forward 5’GACATAGAAGTTACTGGCGATAG3’, SARS-CoV-2_Orf1ab_Reverse 5’TTAATATGACGCGCACTACAG3’, and SARS-CoV-2_Orf1ab_Probe 5’ HEX-ACCCCGTGACCTTGGTGCTTGT-BHQ-1 3’. Hamster β-actin was used as a reference gene using the following primers hACTB-F 5ʹCTCCCAGCACCATGAAGATC3ʹ, hACTB-R 5ʹGCTGGAAGGTGGACAGTG3ʹ, and hACTB-Probe 5ʹ Cy5-TGTGGATCGGTGGCTCCATCCTG-BHQ-3 3ʹ. Normalisation was performed using the ΔΔCt method and SARS-CoV-2 genomic copies were calculated by interpolating adjusted ORF1ab Ct values from a standard curve. All runs were performed on the ABI7500 Fast instrument and results analysed with the 7500 Software v2.3 (Applied Biosystems, Life Technologies).

### Histology, immunohistochemistry and *in situ* hybridization

3-µm thick sections of formalin-fixed (8%) and paraffin-wax embedded (FFPE) tissues (lung, trachea, larynx, and EDTA-decalcified turbinates) were cut and mounted on glass slides. Slides were stained with haematoxylin and eosin (HE). The following antibodies were used: anti-SARS CoV-2 nucleocapsid (Novus Biologicals), anti-CD3 (Agilent DAKO) anti-Pax5 (Agilent DAKO), TTF-1 (Leica), IBA1 (Fujifilm), and MX1 (Cell Signaling). As negative control, the primary antibody was replaced by isotype serum. For visualization, the EnVision+/HRP, Mouse, HRP kit (Agilent DAKO) or EnVision+/HRP, Rabbit, HRP kit (Agilent DAKO), respectively, was used in an automated stainer (Autostainer Link 48, Agilent Technologies). For all immunohistochemistry experiments, 3,3’-Diaminobenzidine (DAB) was used as a chromogen. RNA was detected using RNAScope according to manufacturer’s instructions (Advanced Cell Diagnostics, RNAScope) with simmering in target solution and proteinase K treatment. The following probes were used: SARS-CoV-2 spike (product code: 848561), DapB (product code: 310043), Ubiquitin (product code: 310041) as well as cricetine MX1 (product code: 1153131), IFIT1 (product code: 1153111), OAS1 (product code: 1153101), RSAD2 (product code: 1153121).

### Digital pathology

For image analysis, slides were scanned with an Aperio VERSA 8 Brightfield, Fluorescence & FISH Digital Pathology Scanner (Leica Biosystems) at 200 x brightfield magnification. Areas to be analysed (whole lung, nasal, laryngeal, tracheal as well as upper and lower bronchial epithelium) were manually outlined using QuPath (Version 0.3.2) [63], HALO® (Indica Labs) or Aperio ImageScope (Leica Biosystems). For each set of immunohistochemistry or *in situ*-hybridization, respectively, the algorithm to detect the percentage of positive cells (IBA1, CD3, PAX5) or positively stained area in pixel (MX1) was tuned individually before analysing the slides of one experiment and marker. For the quantification of areas with proliferating type-2-pneumocytes, the whole lung area was outlined. Subsequently, the HALO algorithm “AI” was trained on several slides to detect clusters of TTF1^+^ nuclei within the lung corresponding to alveola epithelial hyperplasia (“medusa”) while ignoring individual TTF1^+^ cells and the positively stained bronchial epithelium. The readout is therefore the percentage of positive TTF-1-stained area in the lungs with medusa shapes. Slides with artefacts were excluded from the analysis as well as artefacts on the slides, which have been manually excluded. All photomicrographs have been captured with Aperio ImageScope (Leica Biosystems).

### Online digital pathology tool

Representative whole scanned images are available online at https://covid-atlas.cvr.gla.ac.uk. The “CVR Virtual Microscope,” is an online tool where users can zoom in and out of digital images of scanned tissues, accessing therefore the same context and information experienced on a microscope.

### RNA sequencing (Bulk RNAseq)

Total RNA extracted from lung and blood samples from uninfected and infected animals was quantified using Qubit Fluorometer 4 (Life Technologies; Q33238), Qubit RNA HS Assay (Life Technologies; Q32855) and dsDNA HS Assay Kits (Life Technologies; Q32854). RNA integrity number was determined on a 4200 TapeStation System (Agilent Technologies; G2991A) using a High Sensitivity RNA Screen Tape assay (Agilent Technologies; 5067–5579). Before library preparation, haemoglobin RNA was removed from 1 µg of RNA extracted from blood samples using the GLOBINclear-Mouse/Rat Globin mRNA Removal Kit (Thermo Fisher Scientific; AM1981) following the manufacturer’s instructions. Total RNA (500 ng) was used to prepare libraries for sequencing using the Illumina TruSeq Stranded mRNA Library Prep kit (Illumina; 20020594) and SuperScript II Reverse Transcriptase (Invitrogen; 18064014) according to the manufacturer’s instructions. The PCR amplified dual indexed libraries were cleaned up with Agencourt AMPure XP magnetic beads (Beckman Coulter; A63881), quantified using Qubit Fluorometer 4 (Life Technologies; Q33238) and Qubit dsDNA HS Assay Kit (Life Technologies; Q32854). Their size distribution was assessed using a 4200 TapeStation System (Agilent Technologies; G2991A) with a High Sensitivity D1000 Screen Tape assay (Agilent Technologies; 5067– 5584). Libraries were pooled in equimolar concentrations and sequenced using high output cartridges with 75 cycles (Illumina; 20024911) on an Illumina NextSeq 550 sequencer (Illumina; SY-415-1002). A Q score of ≥30 was presented in at least 95% of the sequencing reads generated.

### Sequence quality and assembly

Prior to performing bioinformatics analysis, RNA-Seq reads quality was assessed using FastQC software (http://www.bioinformatics.babraham.ac.uk/projects/fastqc). Sequence adaptors were removed using TrimGalore (https://www.bioinformatics.babraham.ac.uk/projects/trim_galore/). Subsequently, the RNA-Seq reads were analysed. Sequence reads were aligned to the *Mesocricetus auratus* genome (MesAur1.0), downloaded via Ensembl using HISAT2. HISAT2 is a fast and sensitive splice-aware mapper, which aligns RNA sequencing reads to mammalian-sized genomes using the FM index strategy [64].

### Differential expression genes analysis

After the alignment to the *Mesocricetus auratus* genome, FeatureCount [65] was used to calculate the mapped reads counts. In this paper, we observed the differential expression genes (DEGs) of mock vs Delta and mock vs BA.1 (2/6 days) on both lung and blood cells. The DESeq2 [66] in Generalized linear models (GLMs) with multi-factor designs (here the factors are gender and condition of the samples) was used for differential expression genes analysis. FDR P-value < 0.05 was used as the cut-off of significant differential expression genes. We analysed the differential expressed gene sets corresponding to molecular pathways of the Reactome database[67].

### Statistical Analysis

All graphs and statistical analyses were produced using GraphPad Prism 7 (GraphPad Software Inc., San Diego, CA, USA) as indicated in each figure legend. P values < 0.05 were deemed to be significant.

### Data Availability Statement

The raw FASTQ files generated during this project have been submitted to the European Nucleotide Archive (ENA) under project accession number PRJEB55782. Raw data underpinning the figures in this study are available in Elighten (doi: XXXXXXX)

## Acknowledgements

We acknowledge the assistance of Catrina Boyd, Scott McCall, and Nicola Munro at Biological Services at the University of Glasgow for guidance on animal experiments, and Lynn Oxford, Lynn Marion Stevenson, Frazer Bell, and Jan Duncan of the Veterinary Pathology Unit excellent technical assistance. We also gratefully acknowledge Diane Vaughan and Alana Hamilton of the III Flow Core Facility at the University of Glasgow for their support & assistance in this work. Funding was provided by the Wellcome Trust (206369/Z/17/Z), the UKRI (to the G2P-UK consortium, MR/W005611/1), by LifeArc (COVID-19 grant) and MRC (MC_UU_00034/5).

## Author’s Contribution

Conceptualization; GRM, VH, AHP, MP; Methodology: GRM, VH, QG, KN, ADSF, WF, JB; Software: QG, DWW; Validation: GRM, VH; Formal analysis: GRM, VH, QG, KN, XH, GI, JA; Investigation: GRM, VH, KK, JA, DCM, XH, SMA, GI, KN, QG, NU, MC; Resources: GDL, VC, JB, WB; Data curation: GRM, VH; Writing-original draft preparation: GRM, VH, AHP, MP; Writing-Review & editing: all; Visualization: GRM, VH, MP; Supervision: GRM, VH, ADSF, WB, AHP, MP; Project administration: GRM, VH, AHP, MP; Funding acquisition: AHP, MP

## Supplementary Data

**Figure S1. Disease Scoring and PCA plots from transcriptomic analysis of tissues from SARS-CoV-2 experimentally infected hamsters.** (A) Hamsters were infected with Delta or BA.1 intranasally. Mock-infected animals received media alone. The weights and disease scores of each animal was recorded daily. (B) All animals were culled 2- or 6-days post-infection (dpi), and RNA was extracted from lungs and blood for RNAseq. PCA plots indicate the variance between the different samples at 2 and 6 dpi. Data were obtained from n=8 animals per group from two independent experiments (4 females and 4 males per group).

**Figure S2. Detection of spike protein or RNA in SARS-CoV-2 experimentally infected hamsters.** Micrographs of lung tissues collected from hamsters infected intranasally with either Delta or BA.1 (or mock-infected). Animals were culled at 2 dpi and lung sections were assessed for the presence of viral protein by immunohistochemistry, or viral RNA by *in situ* hybridisation.

**Figure S3. Tissue Distribution of Delta and BA.1 in experimentally infected hamsters at day 1 post-infection.** (A) Hamsters were infected with either Delta or BA.1 intranasally (or mock infected). Animals were culled 1 day post infection and turbinates, trachea and lungs were collected for digital pathology analyses. (A). Tissues were assessed for the presence of spike RNA by *in situ* hybridisation (B) or for the expression of MX1 by immunohistochemistry (C). For signal quantification, slides were scanned with an Aperio VERSA 8 Brightfield, Fluorescence & FISH Digital Pathology Scanner (Leica Biosystems) at 200 x brightfield magnification. (D) Nasal washes, throat swabs and lungs were analysed for the presence of SARS-CoV-2 genomic RNA by RT-qPCR. Statistical analysis was performed using an unpaired t test, *<0.05, **<0.01, ****<0.0001. Data is representative of two independent experiments, n=6 (3 females and 3 males per group). Blue scale bar: 500 μm; green scale bar: 200 μm. Graphics made using biorender.com.

**Figure S4. Comparison of virulence of SARS-CoV-2 variants used in this study.** Data shown in Fig. 6A, D, E, G and H was merged for either (A) IBA1 or (B) medusas by normalising results to those obtained in BA.1 infected hamsters (taken as 1). Statistical analysis was performed using a One-Way ANOVA with Tukey’s multiple comparisons test. Significance is indicated with *<0.05, **<0.01, ***<0.001, ****<0.0001.

## References

1. Carabelli AM, Peacock TP, Thorne LG, Harvey WT, Hughes J, Peacock SJ, et al. SARS-CoV-2 variant biology: immune escape, transmission and fitness. Nat Rev Microbiol. 2023;21(3):162–77. Epub 2023/01/19. doi: 10.1038/s41579-022-00841-7. PubMed PMID: 36653446; PubMed Central PMCID: PMCPMC9847462.

2. Tracking SARS-CoV-2 Variants. [Internet]. 2022. Available from: https://www.who.int/activities/tracking-SARS-CoV-2-variants (2022).

3. Tegally H, Moir M, Everatt J, Giovanetti M, Scheepers C, Wilkinson E, et al. Emergence of SARS-CoV-2 Omicron lineages BA.4 and BA.5 in South Africa. Nature Medicine. 2022;28(9):1785-90. doi: 10.1038/s41591-022-01911-2.

4. Tegally H, Moir M, Everatt J, Giovanetti M, Scheepers C, Wilkinson E, et al. Emergence of SARS-CoV-2 Omicron lineages BA.4 and BA.5 in South Africa. Nat Med. 2022;28(9):1785-90. Epub 2022/06/28. doi: 10.1038/s41591-022-01911-2. PubMed PMID: 35760080; PubMed Central PMCID: PMCPMC9499863 Lancet Laboratories.

5. Viana R, Moyo S, Amoako DG, Tegally H, Scheepers C, Althaus CL, et al. Rapid epidemic expansion of the SARS-CoV-2 Omicron variant in southern Africa. Nature. 2022;603(7902):679-86. Epub 2022/01/19. doi: 10.1038/s41586-022-04411-y. PubMed PMID: 35042229; PubMed Central PMCID: PMCPMC8942855.

6. Saito A, Tamura T, Zahradnik J, Deguchi S, Tabata K, Anraku Y, et al. Virological characteristics of the SARS-CoV-2 Omicron BA.2.75 variant. Cell Host Microbe. 2022;30(11):1540-55.e15. Epub 2022/10/23. doi: 10.1016/j.chom.2022.10.003. PubMed PMID: 36272413; PubMed Central PMCID: PMCPMC9578327.

7. Ito J, Suzuki R, Uriu K, Itakura Y, Zahradnik J, Kimura KT, et al. Convergent evolution of SARS-CoV-2 Omicron subvariants leading to the emergence of BQ.1.1 variant. Nature Communications. 2023;14(1):2671. doi: 10.1038/s41467-023-38188-z.

8. Tamura T, Ito J, Uriu K, Zahradnik J, Kida I, Anraku Y, et al. Virological characteristics of the SARS-CoV-2 XBB variant derived from recombination of two Omicron subvariants. Nat Commun. 2023;14(1):2800. Epub 2023/05/17. doi: 10.1038/s41467-023-38435-3. PubMed PMID: 37193706; PubMed Central PMCID: PMCPMC10187524 of patents (PCT/JP2016/057254; “Method for inducing differentiation of alveolar epithelial cells”, PCT/JP2016/059786, “Method of producing airway epithelial cells”). The other authors declare that no competing interests exist.

9. Telenti A, Arvin A, Corey L, Corti D, Diamond MS, García-Sastre A, et al. After the pandemic: perspectives on the future trajectory of COVID-19. Nature. 2021;596(7873):495-504. Epub 2021/07/09. doi: 10.1038/s41586-021-03792-w. PubMed PMID: 34237771.

10. Sigal A, Milo R, Jassat W. Estimating disease severity of Omicron and Delta SARS-CoV-2 infections. Nature Reviews Immunology. 2022;22(5):267–9. doi: 10.1038/s41577-022-00720-5.

11. Hyams C, Challen R, Marlow R, Nguyen J, Begier E, Southern J, et al. Severity of Omicron (B.1.1.529) and Delta (B.1.617.2) SARS-CoV-2 infection among hospitalised adults: A prospective cohort study in Bristol, United Kingdom. Lancet Reg Health Eur. 2023;25:100556. Epub 2022/12/20. doi: 10.1016/j.lanepe.2022.100556. PubMed PMID: 36530491; PubMed Central PMCID: PMCPMC9742675.

12. Esper FP, Adhikari TM, Tu ZJ, Cheng Y-W, El-Haddad K, Farkas DH, et al. Alpha to Omicron: Disease Severity and Clinical Outcomes of Major SARS-CoV-2 Variants. The Journal of Infectious Diseases. 2022;227(3):344–52. doi: 10.1093/infdis/jiac411.

13. Pascall DJ, Vink E, Blacow R, Bulteel N, Campbell A, Campbell R, et al. Directions of change in intrinsic case severity across successive SARS-CoV-2 variant waves have been inconsistent. J Infect. 2023;87(2):128–35. Epub 2023/06/04. doi: 10.1016/j.jinf.2023.05.019. PubMed PMID: 37270070; PubMed Central PMCID: PMCPMC10234362.

14. Halfmann PJ, Iida S, Iwatsuki-Horimoto K, Maemura T, Kiso M, Scheaffer SM, et al. SARS-CoV-2 Omicron virus causes attenuated disease in mice and hamsters. Nature. 2022;603(7902):687-92. Epub 2022/01/22. doi: 10.1038/s41586-022-04441-6. PubMed PMID: 35062015; PubMed Central PMCID: PMCPMC8942849

15. Willett BJ, Grove J, MacLean OA, Wilkie C, De Lorenzo G, Furnon W, et al. SARS-CoV-2 Omicron is an immune escape variant with an altered cell entry pathway. Nat Microbiol. 2022;7(8):1161–79. Epub 2022/07/08. doi: 10.1038/s41564-022-01143-7. PubMed PMID: 35798890; PubMed Central PMCID: PMCPMC9352574.

16. Suzuki R, Yamasoba D, Kimura I, Wang L, Kishimoto M, Ito J, et al. Attenuated fusogenicity and pathogenicity of SARS-CoV-2 Omicron variant. Nature. 2022;603(7902):700-5. Epub 2022/02/02. doi: 10.1038/s41586-022-04462-1. PubMed PMID: 35104835; PubMed Central PMCID: PMCPMC8942852.

17. Armando F, Beythien G, Kaiser FK, Allnoch L, Heydemann L, Rosiak M, et al. SARS-CoV-2 Omicron variant causes mild pathology in the upper and lower respiratory tract of hamsters. Nat Commun. 2022;13(1):3519. Epub 2022/06/21. doi: 10.1038/s41467-022-31200-y. PubMed PMID: 35725735; PubMed Central PMCID: PMCPMC9207884.

18. Imai M, Iwatsuki-Horimoto K, Hatta M, Loeber S, Halfmann PJ, Nakajima N, et al. Syrian hamsters as a small animal model for SARS-CoV-2 infection and countermeasure development. Proc Natl Acad Sci U S A. 2020;117(28):16587–95. Epub 2020/06/24. doi: 10.1073/pnas.2009799117. PubMed PMID: 32571934; PubMed Central PMCID: PMCPMC7368255.

19. Chan JF, Zhang AJ, Yuan S, Poon VK, Chan CC, Lee AC, et al. Simulation of the Clinical and Pathological Manifestations of Coronavirus Disease 2019 (COVID-19) in a Golden Syrian Hamster Model: Implications for Disease Pathogenesis and Transmissibility. Clin Infect Dis. 2020;71(9):2428–46. Epub 2020/03/28. doi: 10.1093/cid/ciaa325. PubMed PMID: 32215622; PubMed Central PMCID: PMCPMC7184405.

20. Port JR, Yinda CK, Owusu IO, Holbrook M, Fischer R, Bushmaker T, et al. SARS-CoV-2 disease severity and transmission efficiency is increased for airborne compared to fomite exposure in Syrian hamsters. Nat Commun. 2021;12(1):4985. Epub 2021/08/19. doi: 10.1038/s41467-021-25156-8. PubMed PMID: 34404778; PubMed Central PMCID: PMCPMC8371001.

21. Shuai H, Chan JF-W, Hu B, Chai Y, Yuen TT-T, Yin F, et al. Attenuated replication and pathogenicity of SARS-CoV-2 B.1.1.529 Omicron. Nature. 2022;603(7902):693-9. doi: 10.1038/s41586-022-04442-5.

22. Kimura I, Yamasoba D, Tamura T, Nao N, Suzuki T, Oda Y, et al. Virological characteristics of the SARS-CoV-2 Omicron BA.2 subvariants, including BA.4 and BA.5. Cell. 2022;185(21):3992-4007.e16. Epub 2022/10/06. doi: 10.1016/j.cell.2022.09.018. PubMed PMID: 36198317; PubMed Central PMCID: PMCPMC9472642.

23. Saito A, Irie T, Suzuki R, Maemura T, Nasser H, Uriu K, et al. Enhanced fusogenicity and pathogenicity of SARS-CoV-2 Delta P681R mutation. Nature. 2022;602(7896):300-6. doi: 10.1038/s41586-021-04266-9.

24. Tamura T, Yamasoba D, Oda Y, Ito J, Kamasaki T, Nao N, et al. Comparative pathogenicity of SARS-CoV-2 Omicron subvariants including BA.1, BA.2, and BA.5. Commun Biol. 2023;6(1):772. Epub 2023/07/25. doi: 10.1038/s42003-023-05081-w. PubMed PMID: 37488344; PubMed Central PMCID: PMCPMC10366110 of HiLung, Inc. Y.Y. is a co-inventor of a patent (PCT/JP2016/057254, “Method for inducing differentiation of alveolar epithelial cells”) related to this work. I.Y. reports speaker fees from Chugai Pharmaceutical Co, and AstraZeneca plt, outside the submitted work. The other authors declare no competing interests.

25. Uraki R, Halfmann PJ, Iida S, Yamayoshi S, Furusawa Y, Kiso M, et al. Characterization of SARS-CoV-2 Omicron BA.4 and BA.5 isolates in rodents. Nature. 2022;612(7940):540-5. Epub 2022/11/03. doi: 10.1038/s41586-022-05482-7. PubMed PMID: 36323336.

26. Uraki R, Iida S, Halfmann PJ, Yamayoshi S, Hirata Y, Iwatsuki-Horimoto K, et al. Characterization of SARS-CoV-2 Omicron BA.2.75 clinical isolates. Nature Communications. 2023;14(1):1620. doi: 10.1038/s41467-023-37059-x.

27. Frere JJ, Serafini RA, Pryce KD, Zazhytska M, Oishi K, Golynker I, et al. SARS-CoV-2 infection in hamsters and humans results in lasting and unique systemic perturbations after recovery. Sci Transl Med. 2022;14(664):eabq3059. Epub 2022/07/21. doi: 10.1126/scitranslmed.abq3059. PubMed PMID: 35857629; PubMed Central PMCID: PMCPMC9210449.

28. Chiba S, Kiso M, Nakajima N, Iida S, Maemura T, Kuroda M, et al. Co-administration of Favipiravir and the Remdesivir Metabolite GS-441524 Effectively Reduces SARS-CoV-2 Replication in the Lungs of the Syrian Hamster Model. mBio. 2021;13(1):e0304421. Epub 2022/02/02. doi: 10.1128/mbio.03044-21. PubMed PMID: 35100870; PubMed Central PMCID: PMCPMC8805032.

29. Heydemann L, Ciurkiewicz M, Beythien G, Becker K, Schughart K, Stanelle-Bertram S, et al. Hamster model for post-COVID-19 alveolar regeneration offers an opportunity to understand post-acute sequelae of SARS-CoV-2. Nat Commun. 2023;14(1):3267. Epub 2023/06/06. doi: 10.1038/s41467-023-39049-5. PubMed PMID: 37277327; PubMed Central PMCID: PMCPMC10241385.

30. Liu X, Park HS, Matsuoka Y, Santos C, Yang L, Luongo C, et al. Live-attenuated pediatric parainfluenza vaccine expressing 6P-stabilized SARS-CoV-2 spike protein is protective against SARS-CoV-2 variants in hamsters. PLoS Pathog. 2023;19(6):e1011057. Epub 2023/06/23. doi: 10.1371/journal.ppat.1011057. PubMed PMID: 37352333; PubMed Central PMCID: PMCPMC10325082 number 63/180,534, entitled “Recombinant chimeric bovine/human parainfluenza virus 3 expressing SARS-CoV-2 spike protein and its use”, filed by the United States of America, Department of Health and Human Services.

31. Machado RRG, Walker JL, Scharton D, Rafael GH, Mitchell BM, Reyna RA, et al. Immunogenicity and efficacy of vaccine boosters against SARS-CoV-2 Omicron subvariant BA.5 in male Syrian hamsters. Nat Commun. 2023;14(1):4260. Epub 2023/07/18. doi: 10.1038/s41467-023-40033-2. PubMed PMID: 37460536; PubMed Central PMCID: PMCPMC10352277.

32. Barut GT, Halwe NJ, Taddeo A, Kelly JN, Schön J, Ebert N, et al. The spike gene is a major determinant for the SARS-CoV-2 Omicron-BA.1 phenotype. Nat Commun. 2022;13(1):5929. Epub 2022/10/08. doi: 10.1038/s41467-022-33632-y. PubMed PMID: 36207334; PubMed Central PMCID: PMCPMC9543931.

33. Yen HL, Sit THC, Brackman CJ, Chuk SSY, Gu H, Tam KWS, et al. Transmission of SARS-CoV-2 delta variant (AY.127) from pet hamsters to humans, leading to onward human-to-human transmission: a case study. Lancet. 2022;399(10329):1070-8. Epub 2022/03/14. doi: 10.1016/s0140-6736(22)00326-9. PubMed PMID: 35279259; PubMed Central PMCID: PMCPMC8912929.

34. Twohig KA, Nyberg T, Zaidi A, Thelwall S, Sinnathamby MA, Aliabadi S, et al. Hospital admission and emergency care attendance risk for SARS-CoV-2 delta (B.1.617.2) compared with alpha (B.1.1.7) variants of concern: a cohort study. Lancet Infect Dis. 2022;22(1):35-42. Epub 2021/08/31. doi: 10.1016/s1473-3099(21)00475-8. PubMed PMID: 34461056; PubMed Central PMCID: PMCPMC8397301

35. Abels E, Pantanowitz L, Aeffner F, Zarella MD, van der Laak J, Bui MM, et al. Computational pathology definitions, best practices, and recommendations for regulatory guidance: a white paper from the Digital Pathology Association. J Pathol. 2019;249(3):286–94. Epub 2019/07/30. doi: 10.1002/path.5331. PubMed PMID: 31355445; PubMed Central PMCID: PMCPMC6852275.

36. Cantwell AM, Singh H, Platt M, Yu Y, Lin YH, Ikeno Y, et al. Kinetic Multi-omic Analysis of Responses to SARS-CoV-2 Infection in a Model of Severe COVID-19. J Virol. 2021;95(20):e0101021. Epub 2021/07/29. doi: 10.1128/jvi.01010-21. PubMed PMID: 34319784; PubMed Central PMCID: PMCPMC8475517.

37. Nouailles G, Wyler E, Pennitz P, Postmus D, Vladimirova D, Kazmierski J, et al. Temporal omics analysis in Syrian hamsters unravel cellular effector responses to moderate COVID-19. Nature Communications. 2021;12(1):4869. doi: 10.1038/s41467-021-25030-7.

38. Hoagland DA, Møller R, Uhl SA, Oishi K, Frere J, Golynker I, et al. Leveraging the antiviral type I interferon system as a first line of defense against SARS-CoV-2 pathogenicity. Immunity. 2021;54(3):557–70.e5. Epub 2021/02/13. doi: 10.1016/j.immuni.2021.01.017. PubMed PMID: 33577760; PubMed Central PMCID: PMCPMC7846242.

39. Zarella MD, Bowman D, Aeffner F, Farahani N, Xthona A, Absar SF, et al. A Practical Guide to Whole Slide Imaging: A White Paper From the Digital Pathology Association. Archives of pathology & laboratory medicine. 2019;143(2):222–34. Epub 2018/10/12. doi: 10.5858/arpa.2018-0343-RA. PubMed PMID: 30307746.

40. Shaw AE, Hughes J, Gu Q, Behdenna A, Singer JB, Dennis T, et al. Fundamental properties of the mammalian innate immune system revealed by multispecies comparison of type I interferon responses. PLoS Biol. 2017;15(12):e2004086. doi: 10.1371/journal.pbio.2004086. PubMed PMID: 29253856.

41. Sia SF, Yan LM, Chin AWH, Fung K, Choy KT, Wong AYL, et al. Pathogenesis and transmission of SARS-CoV-2 in golden hamsters. Nature. 2020;583(7818):834-8. Epub 2020/05/15. doi: 10.1038/s41586-020-2342-5. PubMed PMID: 32408338; PubMed Central PMCID: PMCPMC7394720.

42. Muñoz-Fontela C, Dowling WE, Funnell SGP, Gsell PS, Riveros-Balta AX, Albrecht RA, et al. Animal models for COVID-19. Nature. 2020;586(7830):509-15. Epub 2020/09/24. doi: 10.1038/s41586-020-2787-6. PubMed PMID: 32967005; PubMed Central PMCID: PMCPMC8136862.

43. Bruno MD, Bohinski RJ, Huelsman KM, Whitsett JA, Korfhagen TR. Lung cell-specific expression of the murine surfactant protein A (SP-A) gene is mediated by interactions between the SP-A promoter and thyroid transcription factor-1. J Biol Chem. 1995;270(12):6531–6. Epub 1995/03/24. doi: 10.1074/jbc.270.12.6531. PubMed PMID: 7896788.

44. Stahlman MT, Gray ME, Whitsett JA. Expression of thyroid transcription factor-1(TTF-1) in fetal and neonatal human lung. J Histochem Cytochem. 1996;44(7):673–8. Epub 1996/07/01. doi: 10.1177/44.7.8675988. PubMed PMID: 8675988.

45. Saito A, Tamura T, Zahradnik J, Deguchi S, Tabata K, Anraku Y, et al. Virological characteristics of the SARS-CoV-2 Omicron BA.2.75 variant. Cell Host & Microbe. 2022;30(11):1540-55.e15. doi: https://doi.org/10.1016/j.chom.2022.10.003.

46. Tamura T, Ito J, Uriu K, Zahradnik J, Kida I, Nasser H, et al. Virological characteristics of the SARS-CoV-2 XBB variant derived from recombination of two Omicron subvariants. bioRxiv. 2022:2022.12.27.521986. doi: 10.1101/2022.12.27.521986.

47. Uraki R, Kiso M, Iida S, Imai M, Takashita E, Kuroda M, et al. Characterization and antiviral susceptibility of SARS-CoV-2 Omicron BA.2. Nature. 2022;607(7917):119-27. Epub 2022/05/17. doi: 10.1038/s41586-022-04856-1. PubMed PMID: 35576972.

48. Francis ME, Goncin U, Kroeker A, Swan C, Ralph R, Lu Y, et al. SARS-CoV-2 infection in the Syrian hamster model causes inflammation as well as type I interferon dysregulation in both respiratory and non-respiratory tissues including the heart and kidney. PLoS Pathog. 2021;17(7):e1009705. Epub 2021/07/16. doi: 10.1371/journal.ppat.1009705. PubMed PMID: 34265022; PubMed Central PMCID: PMCPMC8282065.

49. Gruber AD, Firsching TC, Trimpert J, Dietert K. Hamster models of COVID-19 pneumonia reviewed: How human can they be? Vet Pathol. 2022;59(4):528–45. Epub 2021/12/04. doi: 10.1177/03009858211057197. PubMed PMID: 34856819; PubMed Central PMCID: PMCPMC9207993.

50. Boudewijns R, Thibaut HJ, Kaptein SJF, Li R, Vergote V, Seldeslachts L, et al. STAT2 signaling restricts viral dissemination but drives severe pneumonia in SARS-CoV-2 infected hamsters. Nat Commun. 2020;11(1):5838. Epub 2020/11/19. doi: 10.1038/s41467-020-19684-y. PubMed PMID: 33203860; PubMed Central PMCID: PMCPMC7672082.

51. Carsana L, Sonzogni A, Nasr A, Rossi RS, Pellegrinelli A, Zerbi P, et al. Pulmonary post-mortem findings in a series of COVID-19 cases from northern Italy: a two-centre descriptive study. Lancet Infect Dis. 2020;20(10):1135–40. Epub 2020/06/12. doi: 10.1016/s1473-3099(20)30434-5. PubMed PMID: 32526193; PubMed Central PMCID: PMCPMC7279758.

52. Tian S, Xiong Y, Liu H, Niu L, Guo J, Liao M, et al. Pathological study of the 2019 novel coronavirus disease (COVID-19) through postmortem core biopsies. Mod Pathol. 2020;33(6):1007–14. Epub 2020/04/16. doi: 10.1038/s41379-020-0536-x. PubMed PMID: 32291399; PubMed Central PMCID: PMCPMC7156231.

53. Polak SB, Van Gool IC, Cohen D, von der Thüsen JH, van Paassen J. A systematic review of pathological findings in COVID-19: a pathophysiological timeline and possible mechanisms of disease progression. Mod Pathol. 2020;33(11):2128–38. Epub 2020/06/24. doi: 10.1038/s41379-020-0603-3. PubMed PMID: 32572155; PubMed Central PMCID: PMCPMC7306927.

54. Barkauskas CE, Cronce MJ, Rackley CR, Bowie EJ, Keene DR, Stripp BR, et al. Type 2 alveolar cells are stem cells in adult lung. J Clin Invest. 2013;123(7):3025–36. Epub 2013/08/08. doi: 10.1172/jci68782. PubMed PMID: 23921127; PubMed Central PMCID: PMCPMC3696553.

55. Barkauskas CE. A Specialized Few Among Many: Identification of a Novel Lung Epithelial Stem Cell Population. Cell Stem Cell. 2020;26(3):295–6. Epub 2020/03/07. doi: 10.1016/j.stem.2020.02.010. PubMed PMID: 32142655; PubMed Central PMCID: PMCPMC7154508.

56. Vaughan AE, Brumwell AN, Xi Y, Gotts JE, Brownfield DG, Treutlein B, et al. Lineage-negative progenitors mobilize to regenerate lung epithelium after major injury. Nature. 2015;517(7536):621-5. Epub 2014/12/24. doi: 10.1038/nature14112. PubMed PMID: 25533958; PubMed Central PMCID: PMCPMC4312207.

57. Kathiriya JJ, Brumwell AN, Jackson JR, Tang X, Chapman HA. Distinct Airway Epithelial Stem Cells Hide among Club Cells but Mobilize to Promote Alveolar Regeneration. Cell Stem Cell. 2020;26(3):346–58.e4. Epub 2020/01/25. doi: 10.1016/j.stem.2019.12.014. PubMed PMID: 31978363; PubMed Central PMCID: PMCPMC7233183.

58. Lin E, Fuda F, Luu HS, Cox AM, Fang F, Feng J, et al. Digital pathology and artificial intelligence as the next chapter in diagnostic hematopathology. Semin Diagn Pathol. 2023;40(2):88–94. Epub 2023/02/22. doi: 10.1053/j.semdp.2023.02.001. PubMed PMID: 36801182.

59. Marletta S, L’Imperio V, Eccher A, Antonini P, Santonicco N, Girolami I, et al. Artificial intelligence-based tools applied to pathological diagnosis of microbiological diseases. Pathol Res Pract. 2023;243:154362. Epub 2023/02/10. doi: 10.1016/j.prp.2023.154362. PubMed PMID: 36758417.

60. Wu B, Moeckel G. Application of digital pathology and machine learning in the liver, kidney and lung diseases. J Pathol Inform. 2023;14:100184. Epub 2023/01/31. doi: 10.1016/j.jpi.2022.100184. PubMed PMID: 36714454; PubMed Central PMCID: PMCPMC9874068.

61. Yamasoba D, Kimura I, Nasser H, Morioka Y, Nao N, Ito J, et al. Virological characteristics of the SARS-CoV-2 Omicron BA.2 spike. Cell. 2022;185(12):2103-15.e19. Epub 2022/05/15. doi: 10.1016/j.cell.2022.04.035. PubMed PMID: 35568035; PubMed Central PMCID: PMCPMC9057982.

62. Rihn SJ, Merits A, Bakshi S, Turnbull ML, Wickenhagen A, Alexander AJT, et al. A plasmid DNA-launched SARS-CoV-2 reverse genetics system and coronavirus toolkit for COVID-19 research. PLoS Biol. 2021;19(2):e3001091. Epub 2021/02/26. doi: 10.1371/journal.pbio.3001091. PubMed PMID: 33630831; PubMed Central PMCID: PMCPMC7906417.

63. Bankhead P, Loughrey MB, Fernández JA, Dombrowski Y, McArt DG, Dunne PD, et al. QuPath: Open source software for digital pathology image analysis. Sci Rep. 2017;7(1):16878. Epub 2017/12/06. doi: 10.1038/s41598-017-17204-5. PubMed PMID: 29203879; PubMed Central PMCID: PMCPMC5715110.

64. Kim D, Langmead B, Salzberg SL. HISAT: a fast spliced aligner with low memory requirements. Nat Methods. 2015;12(4):357–60. Epub 2015/03/10. doi: 10.1038/nmeth.3317. PubMed PMID: 25751142; PubMed Central PMCID: PMCPMC4655817.

65. Liao Y, Smyth GK, Shi W. featureCounts: an efficient general purpose program for assigning sequence reads to genomic features. Bioinformatics. 2014;30(7):923–30. Epub 2013/11/15. doi: 10.1093/bioinformatics/btt656. PubMed PMID: 24227677.

66. Love MI, Huber W, Anders S. Moderated estimation of fold change and dispersion for RNA-seq data with DESeq2. Genome Biol. 2014;15(12):550. Epub 2014/12/18. doi: 10.1186/s13059-014-0550-8. PubMed PMID: 25516281; PubMed Central PMCID: PMCPMC4302049.

67. Fabregat A, Sidiropoulos K, Viteri G, Forner O, Marin-Garcia P, Arnau V, et al. Reactome pathway analysis: a high-performance in-memory approach. BMC Bioinformatics. 2017;18(1):142. Epub 2017/03/03. doi: 10.1186/s12859-017-1559-2. PubMed PMID: 28249561; PubMed Central PMCID: PMCPMC5333408.

